# Heatrich-BS enables efficient CpG enrichment and highly scalable cell-free DNA methylation profiling

**DOI:** 10.1101/2021.04.05.437976

**Authors:** Elsie Cheruba, Ramya Viswanathan, Pui-Mun Wong, Howard John Womersley, Yiting Lau, Anna Gan, Polly S.Y. Poon, Sarah B. Ng, Dawn Qingqing Chong, Iain Beehuat Tan, Lih Feng Cheow

## Abstract

Genome wide analysis of cell-free DNA (cfDNA) methylation profile has been shown to be a promising approach for sensitive and specific multi-cancer detection. However, scaling these assays for clinical translation is impractical due to the high cost of whole genome bisulfite sequencing. We showed that the small fraction of GC-rich genome is highly enriched in CpG sites and disproportionately harbored the majority of cancer-specific methylation signature. Here, we report on the simple but effective Heat enrichment of CpG-rich regions for Bisulfite Sequencing (Heatrich-BS) platform that enables focused methylation profiling in these highly informative regions. Our novel method and bioinformatics algorithm enable high accuracy and sensitivity in tumor burden estimation and quantitative monitoring of colorectal patient response to treatment, at much reduced sequencing requirement. Heatrich-BS holds great potential for highly scalable screening and regular monitoring of cancer using liquid biopsy.

---

Recent studies have demonstrated the promising use of methylation profiling of cell-free DNA (cfDNA) for multi-cancer detection, leveraging on tissue and cancer specific methylation patterns^1,2^. Methylation-profiling of cfDNA has been shown to outperform mutation assays in cancer detection and tissue of origin localization^2^. Such improved detection can be achieved because methylation-based cfDNA assays offer the following advantages: (i) Methylation patterns are specific to the tissue and type of cancer. This enables the use of published data from resected tumors to generate a unique methylation signature for the cancer of interest^3^. (ii) The methylome of cancer cells exhibit a distinct pattern of methylation changes – hypermethylation of CpG islands (CGIs) and hypomethylation of the genome^4^. Despite their advantages, the high cost associated with the high number of sequencing reads required (>100 million reads) for whole-genome cfDNA methylation assays makes them impractical for routine use.

The need to cover sufficient informative regions in the genome necessitates a high number of sequencing reads. Whole genome bisulfite sequencing (WGBS), the most comprehensive and unbiased method to profile the epigenome, requires at least 30 million reads to achieve 1X coverage of the human genome. However, methylation in human DNA predominantly occurs in the CpG context, and the human genome is generally CpG poor, punctuated by short stretches of CpG-rich regions that collocate with many gene regulatory elements such as promoters. DNA methylation differences between tissue and disease is disproportionately found in these CpG-rich regions that form <1% of the genome^4^. Reduced representation bisulfite sequencing (RRBS), which enriches for these regions by selecting short fragments flanked by CG-containing restriction enzyme sites, allows for selective profiling of these information-rich areas. A wealth of public methylation databases of various healthy and diseased tissues have been generated by applying RRBS. However, RRBS is expected to have limited selectivity on cfDNA, as many cfDNA fragments that are not flanked by the restriction sites will also be sequenced. On the other hand, while alternative approaches based on hybridization capture^2^ can focus the sequencing on informative regions, there is a very high upfront cost involved for optimal probe design. Due to permutations of the methylation state in DNA, using completely methylated or completely unmethylated probes renders this method ineffective when heterogeneous methylation is involved^2^. Single nucleotide variations (SNVs), which are especially common in cancer, would also affect capture of tumor-derived fragments using these probes. Furthermore, once a panel is designed, it is inflexible to be applied to other types of cancers that have a different set of differentially methylated regions (DMRs). Currently, there is a lack of a simple, cost-effective method to enrich for epigenetically important regulatory regions based on natural features of genome content in the liquid biopsy context.

Another factor that necessitates high amount of sequencing for detection of tumor fragments in cfDNA is the high depth needed for tumor fraction estimation. Traditionally, the beta value, which is the average methylation of CpG at particular site, is used for estimation of tumor fraction^5–7^. However, tumor contents are typically low (<1%) for early stage cancers, requiring high sequencing depths for reliable detection of tumor fragments. Recently, the analysis of methylation haplotype blocks^1^, which are genomic regions where neighboring CpGs are highly correlated, have open up a new approach for more sensitive detection. Interestingly, these methylation haplotype blocks are highly enriched in CpG-dense regions. Using a new measure of methylation haplotype load (MHL), which considers both the average methylation of all CpGs in the block and also captures patterns of co-methylation on single DNA molecules, the authors are able to improve robustness and sensitivity of detecting tumor-specific cfDNA. In CancerDetector, the authors exploited the co-methylation patterns to assign a tumor and normal probability for each fragment^8^. In this method, a marker-specific tumor fraction estimate is made by considering all reads that fall within the marker region and the tumor fraction from all markers undergoes iterative pruning until their values converges to a final global tumor fraction. Although proven to work at lower sequencing depth for high tumor fraction, more than 2X sequencing depth is still needed to detect low tumor fractions. Altogether, the need for deep sequencing for detecting low tumor fractions is a key cost bottleneck for liquid biopsy.

Therefore, in order to fully utilize the potential of methylation profiling in liquid biopsy, we developed a two-pronged approach. Firstly, we developed a method that effectively enriches for CpG-dense regions, that are known to harbor significant methylation changes in cancer, regardless of their methylation status. This method, known as Heat enrichment of CpG-rich regions for Bisulfite Sequencing (Heatrich-BS), is a novel assay that can enrich for CpG-rich regions using heat denaturation to eliminate low GC-content fragments. While the concept of melting temperature and the relationship between GC content and heat denaturation is well known, this is the first assay that utilizes the relationship between CpG density and GC content to achieve CpG enrichment. The simple workflow of Heatrich-BS makes it very easy to perform and the use of heat denaturation as a selection method is independent of restriction enzyme recognition sequences. Secondly, we developed a new bioinformatics algorithm that assigns a tumor probability for individual fragments, and integrates these fragment-based probabilities across the entire genome to accurately estimate tumor fractions. This is achieved without the need to estimate tumor fractions from any aggregate measurement within individual markers. This approach capitalizes on the widespread epigenetic DMRs, to integrate low coverage measurement in each informative region and allow tumor fraction estimation with much lower sequencing requirement compared to current methods. Together, heat enrichment and genome-wide read aggregation addresses the critical sensitivity and cost issue for implementing cfDNA methylation-based assays for routine cancer screening and monitoring.

## Results

### Enrichment of DMRs in GC-rich regions

The sequence content in the human genome is highly non-uniform. Long stretches of CpG-poor regions are punctuated by short stretches of CpG-dense regions that coincide with important gene regulatory elements such as promoters. These CpG-dense regions are often differentially methylated between tissues and disease such as cancer^1,9,10^. We use DMRfinder^11^ to identify DMRs between publicly available data on DNA methylation in colorectal cancer (CRC) tissue and healthy plasma (details in Step 1 of tumor fraction determination algorithm in Online Methods) and found that nearly 45% of the DMRs lie within CGIs (Fig 1a), which originates from less than 1% of the genome (Fig 1b). As such, it is of value to focus on this small fraction of CpG-rich genome for epigenetic profiling. The traditional approach to enrich for CGIs is RRBS^12^, with a single-tube variant termed single-cell RRBS (scRRBS)^13^ for low input samples. However, this approach is still limited in utility for enrichment of fragmented cfDNA, as very few fragments can meet the requirement for a CCGG cut site on both ends. In a study that uses scRRBS to perform methylation profiling in cfDNA, henceforth referred to as cell-free RRBS, only 6.4% of the reads are located in CGIs^1^. As such, CGI enrichment in fragmented DNA is limited in efficiency.

**Figure 1:**
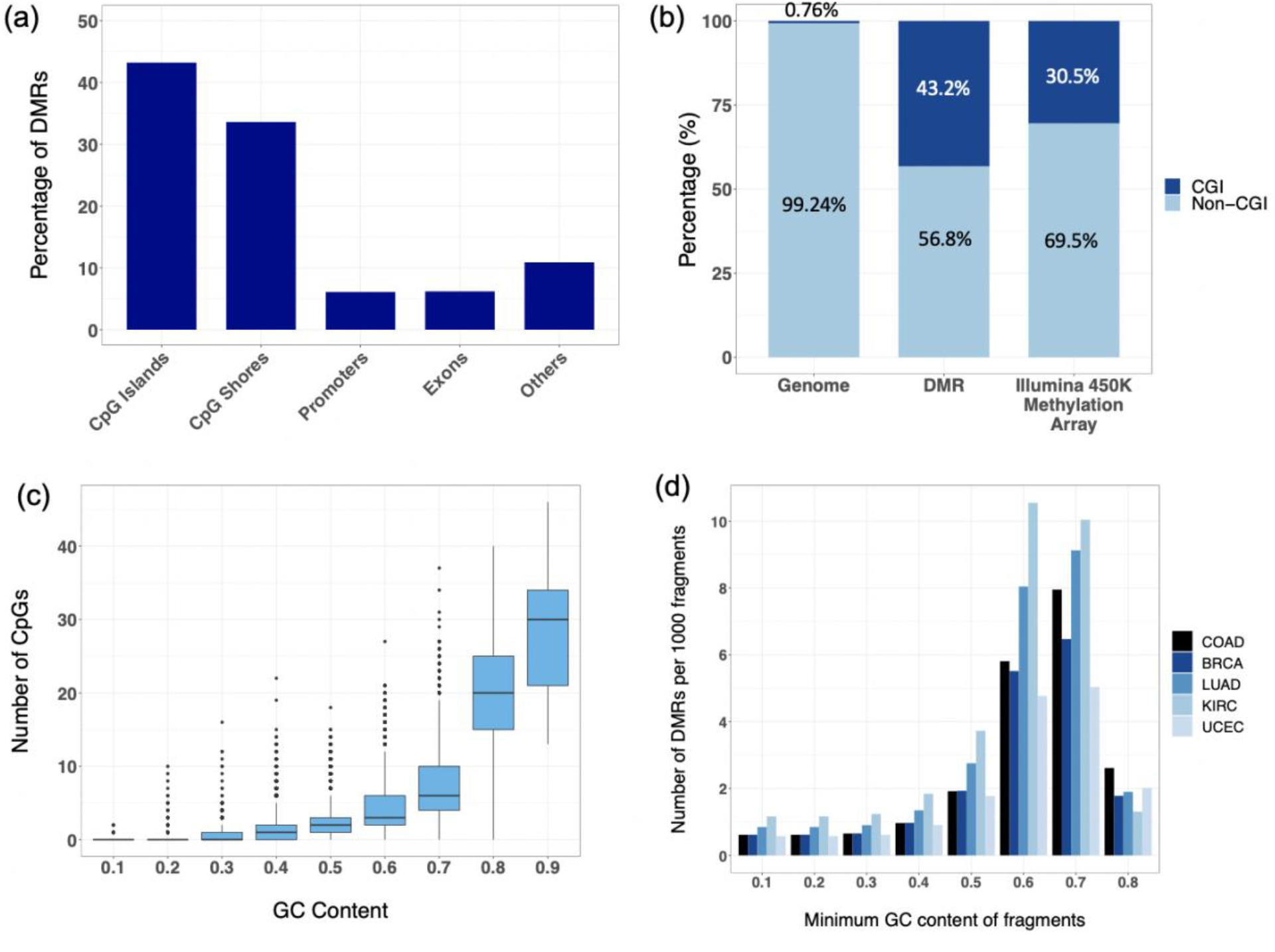
Enrichment of cancer-specific DMRs in CpG-dense and high GC content regions. (a) Percentage of DMRs between colorectal cancer tissue and healthy plasma in different genomic regions. (b) Proportion of DMRs and Illumina 450K Methylation Array probes in CGI with respect to the genomic distribution. (c) Relationship between GC content and number of CpGs in 200bp fragments of the human genome. (d) Number of DMRs of different cancers detected per 1000 fragments using different GC-content thresholds. COAD: colorectal adenocarcinoma, BRCA: breast invasive carcinoma, LUAD: lung adenocarcinoma, KIRC: kidney renal clear cell carcinoma, UCEC: uterine corpus endometrial carcinoma.

While there is no known means to physically enrich for CpG-dense DNA, it is long known that the G+C content of a double-stranded DNA fragment is closely related to its thermal stability. It has been shown that the GC bond in DNA has a binding energy of 25.4 kcal mol^-1^, which is two times stronger than the AT bond, with a binding energy of 12.4 kcal mol^-1^ ^14^. As the presence of a CpG dinucleotide in a fragment adds 2 GC bonds to the duplex, we ask if effective selection of CpG-dense fragments can be achieved by selection of GC-rich fragments. To check this hypothesis, we calculated the GC content and number of CpGs in each fragment in a random sample of 0.5 million 200bp fragments of the human genome (Fig 1c). The excellent correlation between GC content and number of CpGs suggests that CpG-rich fragments can be recovered by selecting DNA fragments with high GC content. Using GC-content as a proxy for CpG-density, we identified that fragments with GC content greater than 0.6 constitutes only 2.5% of the genome, but disproportionately includes 85% of CGIs and 58% of identified DMRs between colorectal cancer tissues and healthy plasma. Therefore, we established that cancer-specific DMRs, which are overrepresented in CpG-dense regions, can be effectively enriched with selection of high-GC DNA fragments.

To further verify if GC-content selection can enable enrichment of cancer-specific DMRs, we identified DMRs between publicly available DNA methylation data of various cancer tissues and healthy plasma. We then determined the number of DMRs that could be detected per 1000 fragments using different GC content thresholds (Fig 1c). Selection of fragments above 0.6 GC content affords nearly 8-fold enrichment of reads in DMRs across different cancers (COAD: colorectal adenocarcinoma, BRCA: breast invasive carcinoma, LUAD: lung adenocarcinoma, KIRC: kidney renal clear cell carcinoma, UCEC: uterine corpus endometrial carcinoma). This enrichment would translate to a reduction is sequencing reads needed to detect tumor-derived fragments as the sequenced fragments are significantly more informative.

### Using heat denaturation to select for GC-rich fragments

Having established the strong correlation between GC content and CpG density, we then explored the use of thermal denaturation as a means of selecting DNA fragments based on GC content. The workflow of the Heatrich-BS assay is shown in Fig 2a. Fragmented DNA was first end repaired and A-tailed. Following this, the sample was heated to denature the GC-poor fragments and adapter ligation was immediately performed. The process of adapter ligation allows selection of intact double-stranded fragments, as T4 DNA ligase has a high selectivity for dsDNA^15^. In this way, only the non-denatured GC-rich DNA fragments are ligated with sequencing adapters. The selected fragments were bisulfite converted and subsequently sequenced.

**Figure 2:**
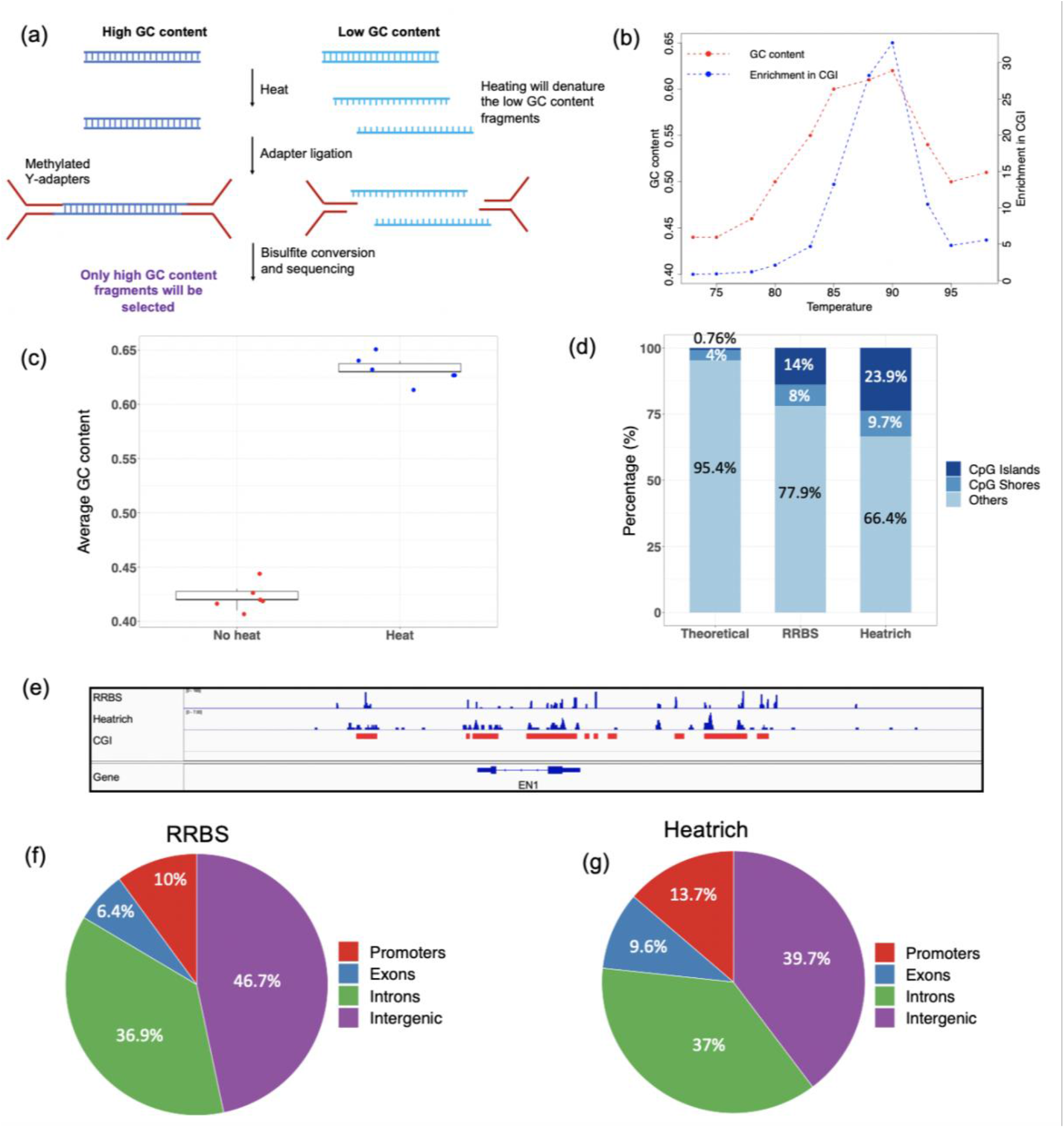
Using heat denaturation to select GC and CpG-rich fragments. (a) Workflow of Heatrich-BS to select for GC-rich fragments. (b) Trend of GC content and read enrichment at CGI over a range of temperatures. (c) Average GC content of sequenced fragments with and without heat denaturation. (d) Distribution of Heatrich and RRBS reads in CpG islands, shores and other regions. (e) Localization of Heatrich and RRBS reads to CGI regions. CGIs are marked in red. (f) Distribution of RRBS reads in different genomic regions. (g) Distribution of Heatrich reads in different genomic regions.

In order to identify the temperature needed to achieve the desired enrichment, we tested a range of temperatures from 75°C to 95°C on sheared genomic DNA (Fig 2b). DNA samples heated to temperatures ranging from 87°C to 90°C had the highest GC content and enrichment of reads in CGI. At higher temperatures, even the fragments with high GC content were denatured, reducing the enrichment in CGIs. We chose 88°C as the optimal denaturation temperature, as this condition had the highest alignment rate along with a high GC content (∼0.62) and enrichment of reads in CGIs (28%). From multiple experiments performed at the selected temperature, we observed that the GC content of the heat-enrichment (Heatrich) samples was much higher than the average GC content of the unheated samples (Fig 2c). The consistent GC content obtained from multiple replicate experiments shows the high degree of reproducibility of our assay.

### Enrichment of CpG-rich regions using Heatrich-BS

Analysis of the mapping of heat-treated sheared DNA showed that there is a significant enrichment of reads that localized to CpG islands and shores, comparing favorably to the theoretical genomic distribution and even RRBS (SRR222486), the gold standard technique for CGI enrichment (Fig 2d). The accumulation of reads around CGIs was visualized (Fig 2e), showing that Heatrich samples displayed significant piling up of reads around CGI. More detailed analysis of the genomic distribution of Heatrich reads showed a remarkable similarity to that of standard RRBS (Fig 2f&g), suggesting that this non-enzymatic approach can be a viable alternative to RRBS for detailed methylation profiling in important genomic regulatory elements. Notably, Heatrich is independent of restriction enzyme sequence, and will thus robustly profile the same regions even in the presence of restriction site polymorphisms, and could find applications when DNA is already fragmented prior to restriction digestion (e.g. FFPE, degraded DNA, cfDNA) where RRBS is not well-suited.

### Heatrich-BS is suitable for CpG-enrichment in cfDNA

cfDNA has promising applications for non-invasive disease detection, and can be suitably profiled using Heatrich-BS due to its fragmented nature. We tested the performance of Heatrich-BS on cfDNA samples obtained from colorectal cancer patients. Bisulfite treatment was performed on samples with and without heat denaturation. We first visualized the reads obtained from Heatrich-BS and compared it with WGBS and previously reported cell-free RRBS^1^, subsampled to similar read counts for comparison (Fig 3a). Mapped reads from cell-free RRBS and WGBS were distributed almost uniformly across all genomic regions. On the other hand, the vast majority of Heatrich-BS reads were concentrated at CpG islands and shores, with little reads in the other regions. This shows the specificity of Heatrich-BS in selecting fragments of high CpG density and high GC content. Furthermore, with comparable read counts, the average height of Heatrich-BS peaks is appreciably more than the other datasets. This shows that Heatrich-BS can obtain higher depth in informative regions using the same number of total sequencing reads.

**Figure 3:**
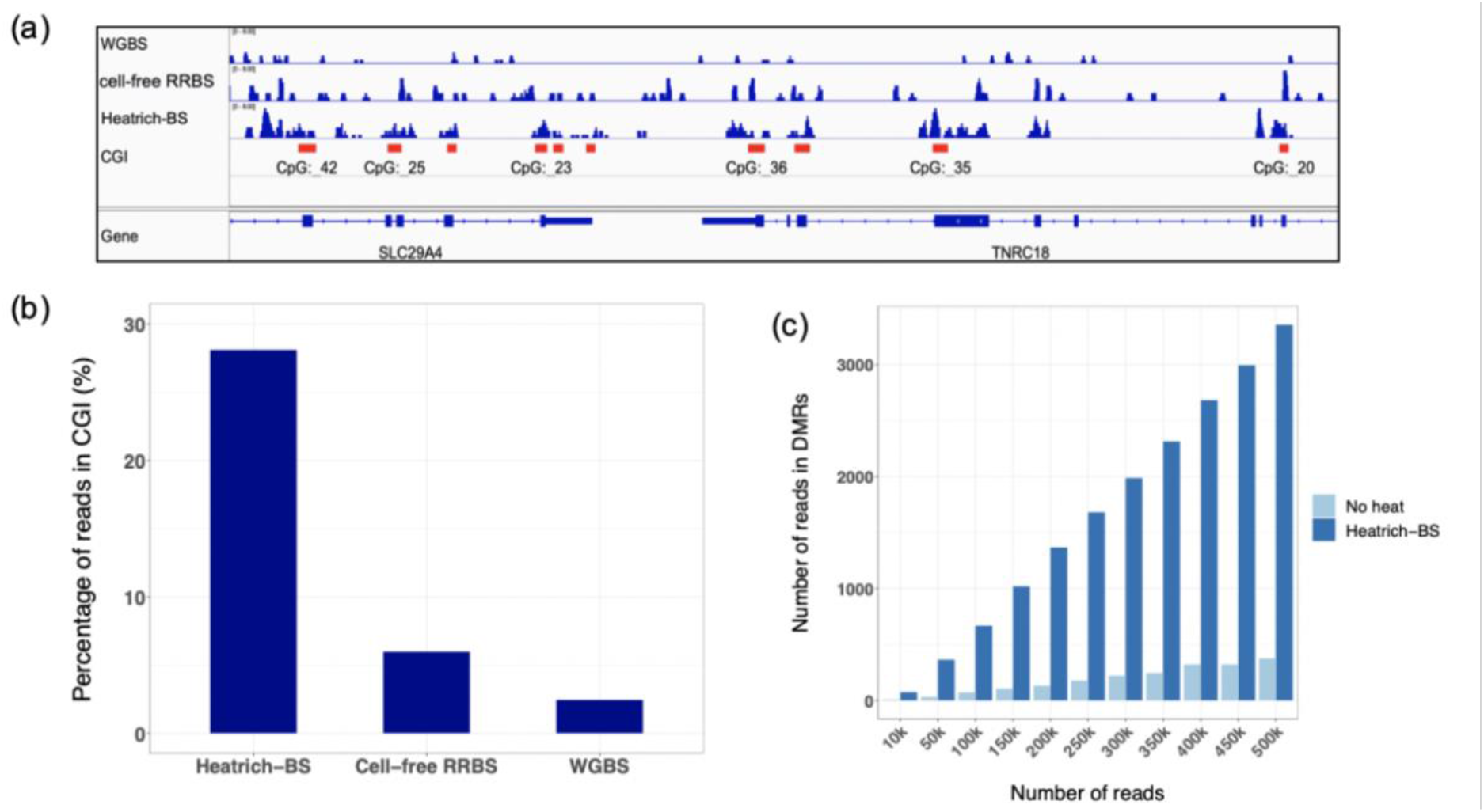
Heatrich-BS effectively enriches cfDNA reads in CpG-rich regions. (a) Piling up of cfDNA reads and localization to CGI using WGBS, RRBS and Heatrich-BS methods. CGI regions are marked in red. (b) Percentage of cfDNA reads in CGI using Heatrich-BS, RRBS and WGBS. (c) Number of cfDNA reads in DMR for different total reads with and without heat denaturation.

The distribution of cfDNA sequenced reads from Heatrich-BS and other methods were quantified, and it was seen that the Heatrich-BS samples displayed up to 15-fold enrichment of reads in CGIs compared to WGBS (Fig 3b). Heatrich-BS also outperformed cell-free RRBS, with 30% of Heatrich-BS reads in CGIs but only 5% of cell-free RRBS reads localized to CGIs. This demonstrates that Heatrich-BS displays more effective CGI enrichment for fragmented DNA, compared to current gold standard methods. Furthermore, the number of reads in DMRs were measured for different number of total sequencing reads (Fig 3c). Heatrich-BS had up to 10-fold more reads localizing to DMRs and Heatrich-BS was able to detect up to 10-fold more DMRs than no heat controls using comparable number of sequencing reads (Supplementary Fig 1). This would provide higher sensitivity in detecting fragments of tumor origin for the same number of sequencing reads. These results highlight the strength of Heatrich-BS, which is the effective enrichment of reads in CGI and DMRs with fewer total sequencing reads.

### Estimating the tumor fraction

Recent studies have shown that the co-methylation patterns within individual DNA fragments can be used to distinguish the origins of cfDNA fragments with higher sensitivity. This has led to development of methods such as those that make use of methylation haplotype load^1^ or α-values^8^ for tumor fraction estimation in cfDNA. However, population-averaged measurements at the marker level are still invariably needed, either as a metric for read discordance within a marker, or as a requisite for removing confounding markers in other study. The need for a minimum read-depth in marker regions, especially for low tumor fraction samples, imposes a bottleneck to further reducing sequencing cost without compromising sensitivity. In this work, we developed a bioinformatics algorithm that estimates global tumor fraction by considering only the tumor probability of individual sequenced fragments, without having to estimate population metrices from individual marker regions. The workflow of the algorithm is shown in Fig 4a and detailed in the online methods. Our developed algorithm allows for accurate estimation of low cfDNA tumor fractions (0.5%) with very low coverage (1X) sequencing data.

**Figure 4:**
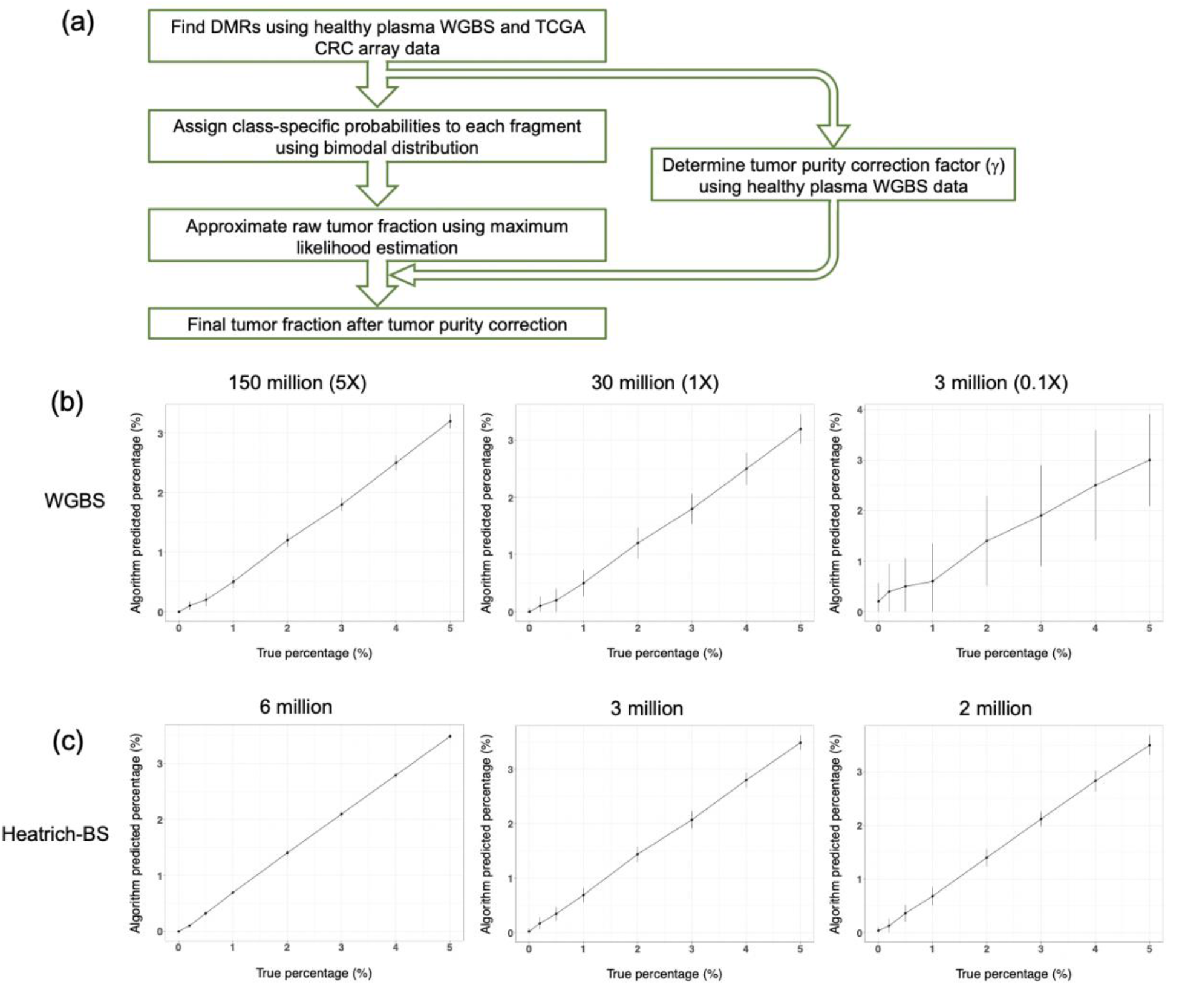
Development and validation of the tumor fraction prediction algorithm. (a) Workflow of tumor fraction prediction algorithm. (b) True and algorithm predicted values of simulated plasma WGBS cfDNA samples at different sequencing depths. (c) True and algorithm predicted value of simulated Heatrich-BS samples using different total sequencing reads.

We first generated a colorectal cancer methylation reference using publicly available Illumina 450K Methylation array data from TCGA, in accordance with previous practices^1,6–8^. We identified DMRs for colorectal cancer using WGBS datasets^5^ of cfDNA from 23 healthy subjects and Illumina 450K Methylation array datasets of 353 colorectal adenocarcinoma samples from TCGA. In order to validate our algorithm with precisely controlled tumor fractions and sequencing depths, we simulated cfDNA of different tumor fractions by mixing WGBS reads (that are not included in reference generation) from plasma of healthy individuals^5^ and tumors of colorectal cancer patients^16^ at different proportions.

When we used the methylation information in the DMRs identified to estimate the tumor fraction of 3 other cfDNA samples from healthy subjects, we observed a non-zero baseline value that was stable regardless of the sequencing depth (Supplementary Fig 2). To identify the source of this baseline, we noted that the resected tumor samples used for generation of methylation data in public databases were often contaminated with non-neoplastic cells. This contamination has been widely acknowledged in previous literature^17,18^ and could lead to overestimation of the tumor fraction in methylation-based analysis. Therefore, we attempted to identify the contribution of these non-neoplastic cells to the colorectal cancer reference, and eliminate its effect on our tumor fraction determination. In order to determine a tumor purity correction factor, we performed Receiver Operating Characteristic (ROC) analysis on simulated plasma cfDNA WGBS samples at 0.5% tumor fraction and identified the correction factor that maximized sensitivity and specificity across multiple sequencing depths (Supplementary Table 1). This correction factor (γ) is dependent on the reference used and only needs to be determined once for each reference.

Using the determined correction factor, we tested our algorithm on simulated plasma WGBS cfDNA samples from 0% to 5% tumor fraction, at different sequencing depths (Fig 4b). At sequencing depths of 5X and 1X, we obtained a high degree of linearity (Pearson correlation >0.99) between the simulated and predicted tumor fraction values while the estimated tumor fraction for the healthy individuals was correctly called as zero. Notably, at 1X depth, where each DMR is covered only once on average, our algorithm is able to accurately detect the presence of small tumor fractions. This is because we can aggregate reads from multiple loci, without requiring high coverage at individual DMRs. Despite this improvement, excessively low coverage would lead to limited number of DMRs being interrogated, which would in turn affect the specificity and confidence of tumor calling. This is seen at very lower depth (0.1X), where we observed larger variations in the predicted tumor fractions, including a higher likelihood of false positives in simulated healthy cfDNA samples. Nevertheless, the sequencing requirement can be kept low without sacrificing coverage at DMRs if the Heatrich-BS assay is used to enrich informative regions. To validate this, we approximated the Heatrich-BS assay by selecting only fragments with GC content more than 0.6 as a simulated plasma cfDNA sample (simulated Heatrich-BS samples). We observed that even using very modest number of total sequencing reads (2-6 million reads), high specificity and tumor calling confidence could be achieved (Fig 4c). Notably, the tumor fraction prediction from simulated Heatrich-BS samples had much higher specificity and lower variance compared to a similar read count simulated WGBS samples (0.1X) at 3 million reads. We also performed ROC analysis to compare the predictive accuracy of using WGBS and Heatrich-BS for low tumor burden detection in cfDNA (0.5% tumor fraction). The predictive accuracy using Heatrich-BS samples is significantly better than conventional WGBS samples (AUC 0.988 vs 0.547) (Supplementary Fig 3). These results demonstrate that Heatrich-BS and the corresponding algorithm enable accurate and confident tumor DNA detection in cfDNA with significantly lesser sequencing requirement compared to existing methods.

### Application of Heatrich-BS on patient cfDNA samples

Finally, we applied Heatrich-BS and the developed algorithm on healthy volunteers and colorectal cancer patient cfDNA samples. We observed that Heatrich-BS reads contain more CpGs per fragment (Supplementary Fig 4), making each sequenced read more informative. Therefore, there were more fragments with tumor probability higher than 90% or lower than 10% when applying the Heatrich-BS method (Supplementary Fig 5), which increases the confidence and accuracy of the tumor fraction called. We then compared our algorithm predicted values to the tumor fraction estimated using the gold standard amplicon sequencing (amplicon-seq) method for mutation detection (Fig 5a). We obtained a Pearson correlation of 0.92 between the Heatrich-BS and amplicon-seq predicted tumor fractions, proving the accuracy of our method and algorithm even in clinical cfDNA samples.

**Figure 5:**
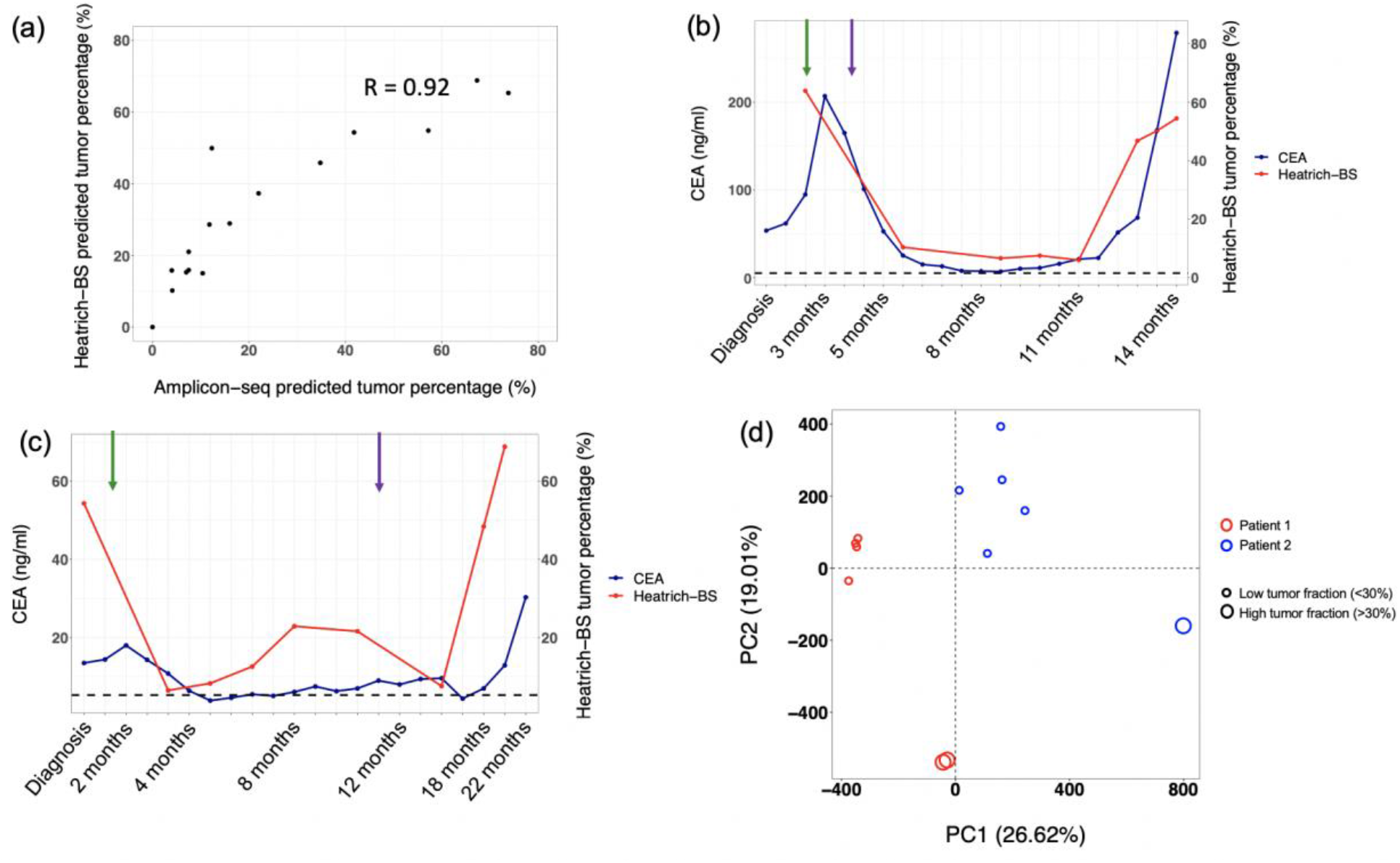
Application of Heatrich-BS on patient cfDNA samples. (a) Amplicon-seq and Heatrich-BS predicted tumor fractions for patient cfDNA samples. (b) Longitudinal tumor monitoring of Patient 1. XELOX (green) and FOLFIRI-Cetuximab (purple) treatment introduction points are marked with arrows. The dashed horizontal line indicates the clinical CEA cut-off. (c) Longitudinal tumor monitoring of Patient 2. 5FU/Oxaliplatin (green) and Irinotecan (purple) treatment introduction points are marked with arrows. The dashed horizontal line indicates the clinical CEA cut-off. (d) PCA analysis of Patients 1 and 2. PC1 separates patients while PC2 separates along tumor burden.

Besides non-invasive cancer diagnosis, liquid biopsy is useful for non-invasive tracking of disease progression. The low cost and high sensitivity for quantitative tumor fraction estimation enabled by Heatrich-BS is attractive for frequent monitoring of patients undergoing treatment and those in remission to detect possibility of relapse. To further validate the applicability of Heatrich-BS in cancer progression monitoring, we tested cfDNA collected from two colorectal cancer patients at different time points during treatment. Using Heatrich-BS, we were able to estimate tumor percentage values and track the patients’ response to treatment (Fig 5b&c). The overall trend in tumor percentage predicted by Heatrich-BS is comparable to the carcinoembryonic antigen (CEA) values, which is a known biomarker for colorectal cancer diagnosis and monitoring. In both patients, the introduction of treatment results in a decline in the tumor fraction, which is seen in Heatrich-BS as well as CEA values, indicating the effectiveness of the treatments administered. However, changes in blood CEA level is not specific to colorectal cancer, and has been shown to only have 34% sensitivity and 84% specificity for recurrence monitoring^19^. Since Heatrich-BS requires fewer reads for tumor burden prediction, and therefore can be performed at lower cost, it could allow more frequent and regular monitoring using this more specific method. Furthermore, CEA level only offers one-dimensional data on the level of cancer while Heatrich-BS can offer multi-dimensional data, including a more quantitative measure of tumor burden as well as other patient-specific information (Fig 5d). It is interesting to note that, on performing Principal Components Analysis (PCA) on the longitudinal patient samples, PC1 separates the patients while PC2 separates along tumor burden. This multi-dimensional information provided by Heatrich-BS can potentially be used to infer sub-type classification, drug resistance and other tumor-specific characteristics. Therefore, cost-effective Heatrich-BS can be used to sensitively monitor tumor progression and characteristics in the clinical context.

## Discussion

The only FDA approved circulating DNA-based assay for colorectal cancer is Epi proColon, which tests for the presence of methylated Septin-9. However, this assay has a sensitivity of only 70% and false positive rate of 20%^20^ as studies have shown that methylated Septin-9 is only present in about 70% of colorectal cancer patients, and Septin-9 methylation is not specific to colorectal cancer^21,22^. Therefore, while such targeted assays are cost-effective and easy to perform, the limited sensitivity and high non-specificity pose challenges for early cancer detection and recurrence monitoring. Genome-wide cfDNA studies like CCGA^2^ use array-based approaches, which can cover more regions, but is also inflexible to incorporate additional informative regions. On the other hand, WGBS would be the most comprehensive method, allowing detection of multiple cancers and subtypes using a single assay. Despite the significant advantages of whole-genome methods, there is no such assay commercially available. This is because WGBS methods are very expensive, making such assays commercially non-viable. Therefore, there is a need for a method to fill the gap between comprehensiveness and cost-effectiveness. In this study, we present a novel assay Heatrich-BS, which uses the well-established concept of melting temperature, to enable CpG enrichment in fragmented DNA. The relationship between GC content and heat denaturation, and the correlation between CpG density and high GC content, was applied in unison to effectively perform enrichment of CpG rich fragments for methylation analysis. This is the first assay that utilizes the well-known concept of melting temperature to achieve CpG enrichment. Heatrich-BS fills the gap between comprehensiveness and cost-effectiveness, by enabling extensive coverage of epigenetically informative regions with low sequencing requirement. It also offers significant advantages compared to targeted methods, as different cancers would not require their own specific panels. The high sensitivity and specificity of Heatrich-BS also allows for its application in early cancer detection. As such, Heatrich-BS can potentially be used as a universal liquid biopsy method for screening and early detection of cancer with high sensitivity, specificity and accuracy.

We have shown that our developed method, Heatrich-BS, was able to enrich for DNA fragments with GC content exceeding 60%. As a result, nearly 30% of Heatrich-BS reads fell into CGIs. This enrichment of reads in CpG-rich regions resulted in greater coverage of DMRs compared to existing methods. We also developed a tumor burden prediction algorithm to augment our assay and validated its application for tumor fractions as low as 0.5%. Furthermore, we have demonstrated proof-of-concept of this novel assay through application on patient cfDNA, enabling deep and comprehensive coverage of cancer-specific DMRs using relatively few total reads. The tumor burden values and trend determined by Heatrich-BS was comparable to the values obtained by orthogonal methods. Therefore, Heatrich-BS and the corresponding algorithm can enable accurate and sensitive prediction of tumor burden in cfDNA.

Heatrich-BS offers significant advantages compared to current assays: (i) The workflow of Heatrich-BS is short and easy to perform. (ii) Heat denaturation is independent of DNA sequence biases that can arise from the use of restriction enzymes in assays like RRBS. This enables its use for CpG enrichment even in genomic DNA, by first performing mechanical fragmentation to obtain short fragments followed by the application of Heatrich-BS. It can also be used to profile fragmented DNA, such as FFPE samples, where current assays could face significant challenges^23^. (iii) Heatrich-BS requires fewer sequencing reads compared to conventional untargeted assays, which makes it cost-effective to perform. This low sequencing depth requirement could prove valuable for applications where targeted methods are limited in scope while genome-wide methods are too expensive.

Given the valuable information that can be obtained from the Heatrich-BS assay, future work can focus on exploring tumor characteristics that can be inferred from the data. Furthermore, since Heatrich-BS data is untargeted, this assay can be extended to multiple cancers, by obtaining multi-cancer DMRs and optimizing the algorithm accordingly. This presents the potential for Heatrich-BS to be a low-cost multi-cancer screening assay, which would be an important innovation to enable the scaling up of cfDNA methylation profiling in liquid biopsy for clinical translation.

## Methods

### Generating sheared DNA

K562 cells (ATCC® CCL-243™) were cultured in high glucose Dulbecco’s modified Eagle’s medium (DMEM) (Gibco) supplemented with 10% Fetal Bovine Serum (FBS) (Gibco) and 1% penicillin-streptomycin (Gibco). Genomic DNA was extracted from cultured K562 cells using the DNeasy Blood and Tissue Kit (Qiagen). The extracted gDNA was fragmented using the LE220 Focused ultrasonicator (Covaris) at the following settings: 450W peak incidence power, 30% duty factor, 200 cycles per burst for 420 sec. The fragmented DNA was size selected for 100-200 bp fragments using BluePippin 2% agarose cassette (Sage Sciences).

### Heatrich-BS protocol

3-5ng of cfDNA was used as input for the Heatrich-BS protocol. Library preparation was done using KAPA Hyper Prep Kit (Kapa Biosystems). 1.4μl of End Repair & A-tailing buffer (Kapa Biosystems) and 0.6μl of End Repair & A-tailing enzyme mix (Kapa Biosystems) was added to 10μl of input DNA, and incubated at 20°C for 30 min, 65°C for 30 min. Following this, the sample was heated at 88°C for 5 min, and immediately placed on ice. The sample was then topped up with 6μl of Ligation buffer (Kapa Biosystems), 2μl of DNA Ligase (Kapa Biosystems), 1μl of nuclease-free water and 1μl of 750nM methylated Truseq adapter (Illumina). For no-heat controls, 1μl of 1.5μM methylated Truseq adapter (Illumina) was used instead. After adding these reagents, the sample was incubated at 25°C for 1 hour and then cleaned up by performing two rounds of 1.2x SPRI Select (Beckman Coulter). The sample was then subject to bisulfite conversion following the recommended protocol of Zymo EZ DNA Methylation kit (Zymo Research). The bisulfite converted DNA was amplified for 15 cycles using Pfu Polymerase (Agilent), cleaned up using 1.2x SPRI Select (Beckman Coulter) and re-amplified using KAPA Hyper Hot-Start Polymerase (Kapa Biosystems) until plateau was reached. The amplified sample was cleaned up using 1.2x SPRI Select (Beckman Coulter), size selected for 190-350 bp fragments using 2% agarose Bluepippin kits (Sage Sciences), quantified using Kapa Library quantification kits (Kapa Biosystems) and sequenced using MiSeq v3 150 cycle kit (Illumina). Pair end sequencing of 75bp each was performed.

### Heatrich-BS Analysis Pipeline

Fastqc^24^ was used to check the quality of the pair-end reads generated by MiSeq. After adapter trimming using Cutadapt^25^, the reads were aligned to the hg38 human genome using Bismark^26^. The aligned reads were deduplicated using Picard tools^27^, following which the Bismark methylation extractor was used to obtain per-base methylation status of each fragment.

### GC content calculation

To calculate the GC content of each fragment, the forward and reverse reads were aligned separately, and then combined to generate a single coordinate range encompassing the entire fragment. The coordinates of the fragment were then used to obtain its sequence from the reference genome. For each fragment, the GC content was defined as the number of Cs and Gs, divided by the total length of the fragment.

### Tumor fraction determination algorithm

The tumor fraction determination algorithm has three major steps:

### Step 1: Identifying the differentially methylated clusters

To identify differentially methylated clusters for tumor-specific cfDNA detection, normal plasma whole genome methylation data^5^ and colorectal adenocarcinoma (COAD) methylation array from TCGA was used. 23 WGBS datasets for normal plasma and 353 Illumina 450K Methylation array datasets from TCGA was used for cluster generation. The TCGA methylation values were extrapolated to ± 100bp of each probe site. To ensure selection of only consistent sites, only methylation values with a standard deviation less than 0.4 between the various sample in that class were chosen, to ensure confidence for the reference. DMRfinder^11^ was used to identify differentially methylated clusters. Within these clusters, sites with a 0.5 difference in methylation were selected.

### Step 2: Using a bimodal distribution to calculate the class-specific probability of each site

Using the generated reference, a normal and tumor class-specific must be assigned to each assayed fragment. Since methylation values are binary, the average methylation value observed in the reference is a proportional combination of the unmethylated and methylated reads. As such, a bimodal distribution can accurately represent the proportional methylation status of the reference. For every site in the reference, the contribution from the unmethylated and methylated mode (0 and 1) was calculated. The relative contributions of each mode in the two classes was used to assign class-specific probabilities for the methylation values in the assayed fragment. In this way, a bimodal reference was used to assign normal or tumor probability values to each site assayed.

### Step 3: Using maximum likelihood estimation to predict the tumor fraction of the sample

After assigning class-specific probabilities to each fragment, the fraction of fragments that come from the tumor must be enumerated. The tumor-derived cfDNA in a sample, also known as tumor fraction, can be denoted as *θ*, where 0 ≤ *θ* < 1. To estimate the tumor fraction *θ*, a maximum likelihood estimation approach and grid search, adapted from CancerDetector^8^, was used to calculate the raw tumor fraction for each sample. The determined tumor purity correction factor (γ) is then applied to the raw tumor fraction to generate the final tumor fraction.

### Data sources

In order to compare our assay with existing methods, we obtained the following data from NCBI GEO: RRBS (SRR222486), cell-free RRBS (GSM2090507). cfDNA WGBS data for algorithm development and validation was obtained by request from the EGA database (EGAS00001001219). Colorectal cancer tumor WGBS data was obtained from NCBI GEO (SRR1035745).

## Supplementary information

Supplementary information is available for this manuscript.

## Acknowledgements

We thank Owen John Llewellyn Rackham for valuable discussions on the algorithm. This work was supported by the Precision medicine and personalised therapeutics seed grant from iHealthtech. Patient samples provided were part of the CaLiBRE program, supported by A*STAR under its IAF-PP scheme (Grant ID: H1801a0019).

## Author contributions

E.C and L.F.C conceived and designed the study; H.J.W performed the pilot experiment; E.C performed the remaining experiments and bioinformatics analysis, with inputs from R.V; P.S.Y.P and S.B.N performed mutation profiling. P.M.W, Y.L, A.G, D.Q.C and I.B.T provided patient samples and inputs on clinical aspects; E.C and L.F.C wrote the manuscript. All authors commented on the manuscript and approved the submission.

## Data availability

The Heatrich-BS next-generation sequencing data for patient samples that support the findings of this study are available upon request from the corresponding author to comply with institutional ethics regulation.

