## Supplemental Figures for "Heatrich-BS enables efficient CpG enrichment and highly scalable cell-free DNA methylation profiling"

### Supplementary Figures

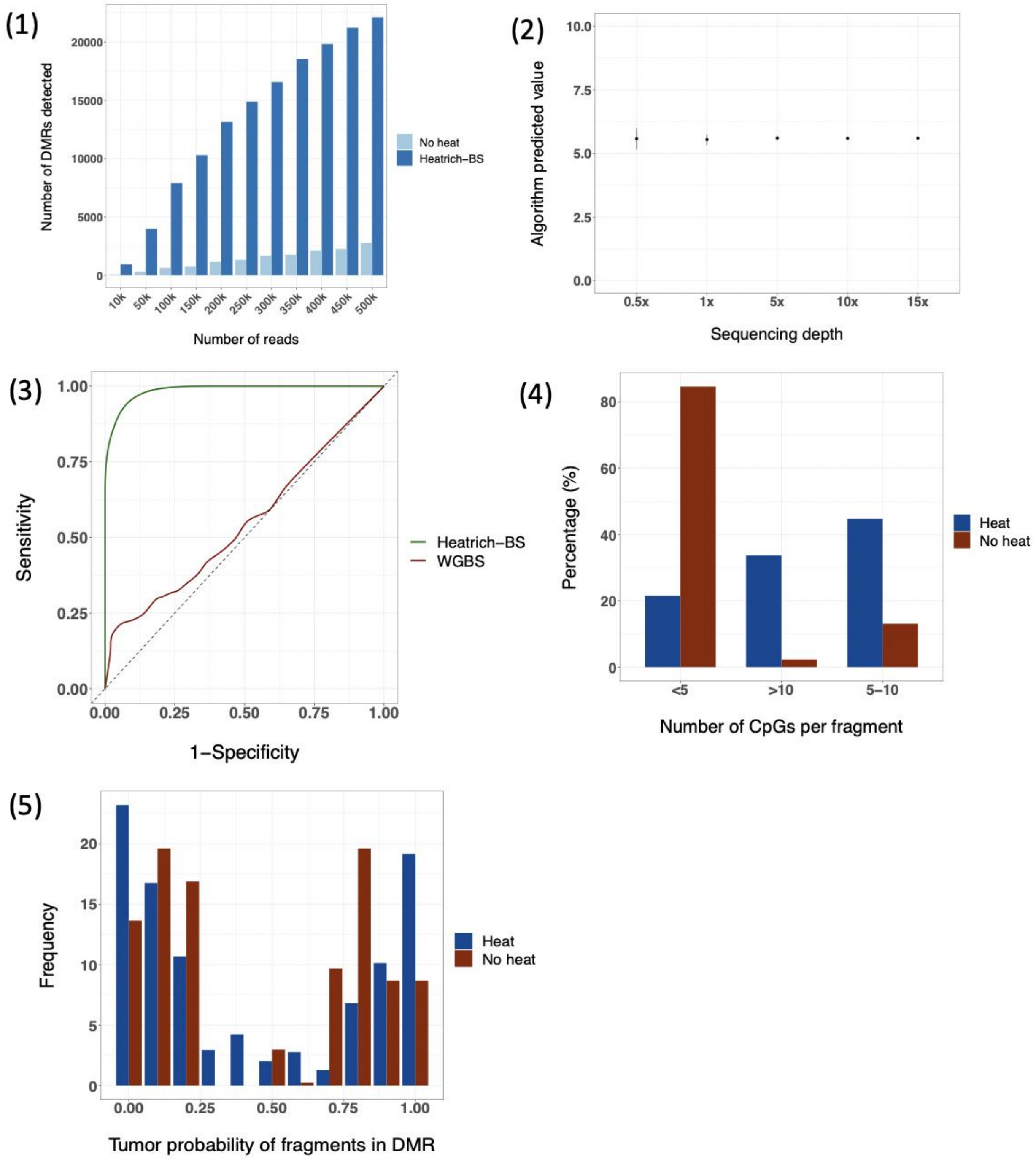


(1) Number of DMRs detected for different total reads with and without heat denaturation. (2) Baseline tumor fraction of normal samples at different sequencing depths. (3) ROC analysis of 0.5% tumor fraction using simulated Heatrich-BS and WGBS cfDNA samples at 3 million total reads. (4) Number of CpGs per fragment with and without heat denaturation. (5) Tumor probability of circulating DNA fragments with and without heat denaturation.

### Supplementary Table

| Threshold \ Depth | 0.5x | | 1x | | 5x | | 10x | | 15x | |
|---|---|---|---|---|---|---|---|---|---|---|
| | 1-Specificity | Sensitivity | 1-Specificity | Sensitivity | 1-Specificity | Sensitivity | 1-Specificity | Sensitivity | 1-Specificity | Sensitivity |
| **0.05** | 0.9 | 1 | 1 | 1 | 1 | 1 | 1 | 1 | 1 | 1 |
| **0.051** | 0.86 | 0.98 | 0.96 | 1 | 1 | 1 | 1 | 1 | 1 | 1 |
| **0.052** | 0.76 | 0.98 | 0.9 | 1 | 1 | 1 | 1 | 1 | 1 | 1 |
| **0.053** | 0.72 | 0.96 | 0.78 | 1 | 1 | 1 | 1 | 1 | 1 | 1 |
| **0.054** | 0.62 | 0.88 | 0.68 | 1 | 0.92 | 1 | 0.98 | 1 | 1 | 1 |
| **0.055** | 0.54 | 0.86 | 0.5 | 0.86 | 0.76 | 1 | 0.74 | 1 | 0.98 | 1 |
| **0.056** | 0.42 | 0.8 | 0.34 | 0.82 | 0.3 | 0.98 | 0.18 | 1 | 0 | 1 |
| **0.057** | 0.38 | 0.64 | 0.18 | 0.76 | 0.02 | 0.96 | 0 | 1 | 0 | 1 |
| **0.058** | 0.22 | 0.56 | 0.06 | 0.64 | 0 | 0.72 | 0 | 0.86 | 0 | 1 |
| **0.059** | 0.14 | 0.46 | 0.02 | 0.44 | 0 | 0.4 | 0 | 0.34 | 0 | 0.02 |
| **0.06** | 0.08 | 0.4 | 0 | 0.38 | 0 | 0.14 | 0 | 0 | 0 | 0 |
| **0.061** | 0.06 | 0.34 | 0 | 0.28 | 0 | 0.02 | 0 | 0 | 0 | 0 |
| **0.062** | 0.04 | 0.32 | 0 | 0.12 | 0 | 0 | 0 | 0 | 0 | 0 |
| **0.063** | 0.04 | 0.24 | 0 | 0.1 | 0 | 0 | 0 | 0 | 0 | 0 |
| **0.064** | 0.02 | 0.2 | 0 | 0.06 | 0 | 0 | 0 | 0 | 0 | 0 |
| **0.065** | 0.02 | 0.12 | 0 | 0.02 | 0 | 0 | 0 | 0 | 0 | 0 |
| **0.066** | 0.02 | 0.04 | 0 | 0 | 0 | 0 | 0 | 0 | 0 | 0 |
| **0.067** | 0.02 | 0 | 0 | 0 | 0 | 0 | 0 | 0 | 0 | 0 |
| **0.068** | 0 | 0 | 0 | 0 | 0 | 0 | 0 | 0 | 0 | 0 |
| **0.069** | 0 | 0 | 0 | 0 | 0 | 0 | 0 | 0 | 0 | 0 |

(1) ROC analysis for different baseline values. The selected threshold with highest sensitivity and specificity is highlighted.
